# Visualizing sounds: training-induced plasticity with a visual-to-auditory conversion device

**DOI:** 10.1101/2021.01.14.426668

**Authors:** Jacques Pesnot Lerousseau, Gabriel Arnold, Malika Auvray

**Author notes:** Co-first authors (Jacques Pesnot Lerousseau, Gabriel Arnold). **Corresponding author:** Malika Auvray, Sorbonne Université, CNRS UMR 7222, Institut des Systèmes Intelligents et de Robotique (ISIR), F-75005 Paris, France, Mail.

## Abstract

Sensory substitution devices aim at restoring visual functions by converting visual information into auditory or tactile stimuli. Although these devices show promises in the range of behavioral abilities they allow, the processes underlying their use remains underspecified. In particular, while an initial debate focused on the visual versus auditory or tactile nature of sensory substitution, since over a decade, the idea that it reflects a mixture of both has emerged. In order to investigate behaviorally the extent to which visual and auditory processes are involved, participants completed a Stroop-like crossmodal interference paradigm before and after being trained with a conversion device which translates visual images into sounds. In addition, participants’ auditory abilities and their phenomenologies were measured. Our study revealed that, after training, when asked to identify sounds, processes shared with vision were involved, as participants’ performance in sound identification was influenced by the simultaneously presented visual distractors. In addition, participants’ performance during training and their associated phenomenology depended on their auditory abilities, revealing that processing finds its roots in the input sensory modality. Our results pave the way for improving the design and learning of these devices by taking into account inter-individual differences in auditory and visual perceptual strategies.

**Highlights:**

- Trained people spontaneously use processes shared with vision when hearing sounds from the device
- Processes with conversion devices find roots both in vision and audition
- Training with a visual-to-auditory conversion device induces perceptual plasticity

## Introduction

Visual-to-auditory and visual-to-tactile sensory substitution devices allow one to transmit visual information by means of the auditory or tactile sensory modality (see Auvray & Myin, 2009, for a review). These devices build on brain plasticity and they are designed with the aim of rehabilitating visual impairments. Most sensory substitution devices consist of a tiny camera, either embedded in glasses or hand-held, recording the external scene in real time. This scene is then converted into sounds or tactile stimuli, with diverse translation codes. In most visual-to-tactile devices, the translation code is analogical. For instance, a visual circle is translated into a circular pattern of tactile stimuli in the “TVSS” device (Bach-y-Rita *et al.*, 1969). Non-analogical codes have also been used, for instance in order to convert distances into vibrations (Maidenbaum *et al.*, 2014, O’Brien, Auvray, Hayward, 2015). The translation code used in visual-to-auditory devices translates several dimensions of the visual signal into dimensions of the auditory signal varying in frequency, loudness, time scanning, or timbre (The vOICe, Meijer, 1992; the Vibe, Hanneton et al., 2010; the EyeMusic, Levy-Tzedek *et al.*, 2014). Behavioural studies have investigated performance across a variety of tasks: sensory substitution devices allow their users to perform localisation tasks (Levy-Tzedek *et al.*, 2012; Proulx *et al.*, 2008), shape recognition (Auvray *et al.*, 2007a, 2007b; Pollok *et al.*, 2005; Striem-Amit *et al.*, 2012b), and navigation tasks (Chebat *et al.*, 2011; 2015).

Beyond this applied purpose, sensory substitution devices allow researchers to investigate the extent to which people can be provided with something resembling vision by means of touch or audition. At first, two main theses have been put forward: the sensory-dominance thesis, according to which perception with a sensory substitution device remains auditory or tactile (Keeley, 2002), and the sensory-deference thesis, according to which perception becomes visual (Bach-y-Rita *et al.*, 2003). For instance, Bach-y-Rita, the pioneer designer for such conversion devices, claimed that their use would allow blind people to see with their skin, or with their brain (Bach-y-Rita *et al.*, 2003).

Early neuroimaging studies did not allow disentangling the question. Supporting the ‘visual’ hypothesis, studies have reported activation of visual occipital areas when trained users of visual-to-auditory devices hear the converted sounds (Poirier, De Volder, & Scheiber, 2007). However, the fact that visual brain areas are activated in response to non-visual stimuli does not necessarily entail that visual images are formed. For instance, although activation increased in visual areas when using a visual-to-tactile device, such activation could underlie tactile functions. Indeed, TMS applied over the visual cortex of blind persons trained with the device results, in some instances, in tactile sensations, and no visual sensations are reported (Kupers *et al.*, 2006). One study directly compared activation longitudinally (Kim & Zatorre, 2011) and did not reveal increased activation, but a change in functional connectivity. This result suggest that the observed cross-modal recruitment of the visual cortex during the use of sensory substitution devices can be due to unmasking of existing computations through non-visual inputs already there prior to devices’ use (Auvray, 2019, Ptito *et al.*, 2005; Striem-Amit *et al.*, 2012; see also Ptito *et al.*, 2018, for a review).

At the functional level, studies have reported some analogies between genuine visual processes and perception with a conversion device. For instance, participants trained with a visual-to-auditory conversion device are able to recognize objects independently of their size, position, or orientation (Kim & Zatorre, 2008). This shows object constancy, as happens during visual object recognition. Trained participants have also been reported to be sensitive to visual illusions, such as the Ponzo illusion (Renier *et al.*, 2005) or the vertical-horizontal illusion (Renier *et al.*, 2006). This gives first indication that some visual mental imagery is involved when using the conversion device.

To overcome the dominance versus deference debate, since over a decade, several researchers advocated that the plasticity at stake in sensory substitution is complex, and based on a multisensory architecture. For instance, the vertical integration thesis (Arnold *et al.*, 2017) suggests that perception with a conversion device goes beyond assimilation to a unique sensory modality (see also Deroy & Auvray, 2012, 2014, for critics of the dominance versus deference alternative). Rather, the processes involved are understood as being vertically integrated within pre-existing capacities that encompass, among other processes, both the substituting and the substituted sensory modalities. Thus, the stimuli coming from the conversion device would activate both visual and auditory brain areas, at low-level, unisensory areas (i.e., visual and auditory primary cortices) or, at higher-level, multisensory areas. In a similar vein, the complementary metamodal/supramodal view of sensory substitution processing (Cecchetti *et al.*, 2016; Heimler, Striem-Amit, & Amedi, 2015; Proulx *et al.*, 2014, 2016) proposes that the functional plasticity at stake occurs at higher levels, involving supramodal representations of shapes with a spatial integration of visual and auditory stimuli. For instance, according to Proulx *et al.* (2014), learning to use a conversion device consists of changing the nature and the complexity of the processes involved. At first, the processes involved would be purely auditory, low level, and very specific. After several hours of training, they would become multisensory, high level, and less specific.

The multisensory view finds support in more recent neuroimaging studies showing that the same brain areas are activated during a task with a visual-to-auditory device and the same task performed visually. For example, the visual extrastriate body-selective area, which strongly responds to images of human bodies, is activated when congenitally blind participants are required to recognize human silhouettes with The vOICe (Striem-Amit & Amedi, 2014). Similarly, recognizing letter with The vOICe has been shown to activate the visual word form area (VWFA) in congenitally blind participants (Striem-Amit *et al.*, 2012). These results led this group to propose that the brain has a flexible task-selective, sensory-independent, supramodal organization rather than a sensory-specific one (Amedi *et al.*, 2017; Heimler *et al.*, 2015).

Although neuroimaging studies point toward a multisensory view, research is still lacking behavioural data to characterize the underlying processes. In addition, the multisensory view also predicts that performance relies not only on supramodal mechanisms but also on the visual and auditory sensory modalities. This involves individual differences as a function of participants’ sensory abilities, which can be investigated only by behavioural methods. As a consequence, the aim of the study reported here is to investigate the multisensory view at the functional level. To do so, the participants were trained with the visual-to-auditory sensory substitution device The vOICe while various behavioural and phenomenological measures were collected. First, the study took advantage of a Stroop-like paradigm (Stroop, 1935) to investigate whether processes shared with vision are involved when using the sensory substitution device. In the original Stroop task, people are requested to name the colour of coloured words. When the meaning of the word is also a colour, it interferes with the participant’s recognition of the colour. For instance, it takes longer to say that the colour of the presented word is blue when the written word reads “red” rather than “blue”. Thus, even if the task consists in naming the colour of a word, reading processes and the ensuing access to the words’ meaning are automatically triggered. The same rationale was used in our study. Both before and after training with The vOICe, the participants completed a Stroop-like task consisting of a sound recognition task with a simultaneous presentation of visual stimuli corresponding to the auditory conversion of visual lines with The vOICe. After training, if processes shared with vision are automatically triggered upon hearing the sounds, for instance if auditory stimuli are mentally converted into visual images or if both stimuli are converted in a common format, then the presentation of visual distractors should interfere with the participants’ performance in the sound recognition task. Thus, processes shared with vision are expected to be revealed with the visual interference paradigm. On the other hand participants’ performance with the device and their phenomenology, are expected to rely on both vision and audition. In order to investigate this, participants’ performance during training was measured. In addition, the participants completed phenomenological questionnaires investigating their qualitative experience with the device as well as low-level auditory tests to assess their auditory abilities. These additional measures allow us to investigate the hypothesis that low-level auditory abilities will be associated with better use of the device, and influence the associated phenomenology.

## Methods

### Participants

Thirty-two naive participants (16 males, 16 females, mean age = 24.1 years, range = 18-35 years) completed the experiment. Half of the participants trained with The vOICe (experimental group). As a control group, the other half did not train with the device. None of the participants were familiar with the device before participating in the study. The participants were assigned randomly to the experimental (N = 16) or to the control (N = 16) group. All of them reported normal or corrected-to-normal vision and normal audition. The study took approximately five hours to complete for the experimental group, and two hours for the control group. The participants received 10 Euros per hour in return for their participation.The experiment was approved by the Local Ethical Committee of the University (“Conseil d’Évaluation Éthique pour les Recherches en Santé”, CERES) it was performed in accordance with the ethical standards laid down in the 1991 Declaration of Helsinki. The participants provided their written informed consent prior to the beginning of the study. Retrospective observed power analysis was based on 10000 Monte Carlo simulations, with the R package ‘simr’ (Green & MacLeod, 2015). The power for the predictor Similarity in the second session of the experimental group, which is our main result, was 78.09% (95% CI: [75.27, 81.90]), for an observed fixed effect of −0.65 ± 0.24. As 80% is the standard power accepted in experimental psychology, we can conclude that our sample size is appropriate.

### Apparatus

The presentation of the stimuli and the recording of data were controlled by a Dell Precision T3400 computer. Images were projected on a 24” Dell P2414H screen. Sounds were transmitted by means of a Sennheiser AKG-272 HD headphone. The sound’s intensity was measured prior to the experiment by means of a Rion NL-42 sonometer. Auditory discrimination tests were conducted with the MatLab toolbox Psychoacoustics (Grassi & Soranzo, 2009). The software Digivox 1.0 was used for the HINT. The device The vOICe (Meijer, 1992) was used to convert the images captured by the camera into sounds (see Figure 1A). During the training phase with The vOICe, the participants were equipped with a camera embedded in a custom-made 3D-printed pair of glasses.

**Figure 1.**
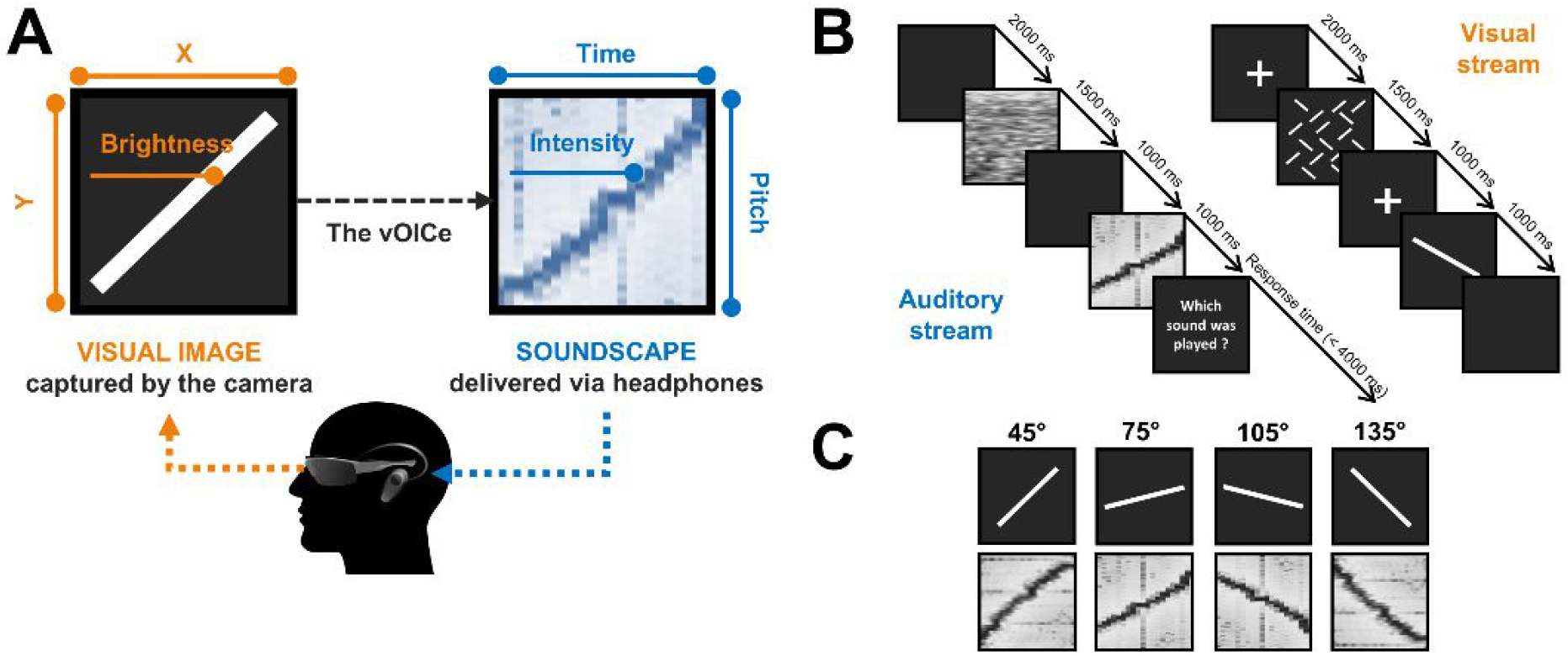
Illustration of the methods. **A.** The visual-to-auditory conversion device (The vOICe). The images recorded by the video camera embedded in the glasses are converted in real-time into soundscapes that are presented to the participants via headphones. The device converts the vertical dimension into auditory frequency, the horizontal dimension into time scanning, and visual brightness into auditory loudness. **B.** Description of the crossmodal interference task. **C.** The auditory targets (spectrogram) and visual distractors that are used in the crossmodal interference task.

### Experimental protocol

The experiment was conducted in an anechoic experimental room. The experiment was divided into three phases: a pre-training phase, a training phase (for the experimental group) or a time interval of similar duration (for the control group), and a post-training phase. The participants completed the auditory tests and the crossmodal interference task during both the pre-training and post-training phases. These phases were strictly equivalent in terms of stimuli, order of presentation, and tests. The participants first completed the auditory tests in the following order: intensity discrimination, pitch discrimination, duration discrimination, and the HINT (Nilsson, Soli, & Sullivan, 1994; Vaillancourt *et al.*, 2005). Then, the participants completed the crossmodal interference task. The time interval between pre-training and post-training sessions was similar for the two groups (experimental group: 2.5 days ± 0.6; control group: 2.4 days ± 0.6; independent t-test: *p* = 0.91). During this interval, the experimental group took part in a 3-hour individual training phase with The vOICe (see Figure 2A). The control group did not participate in this training phase.

**Figure 2.**
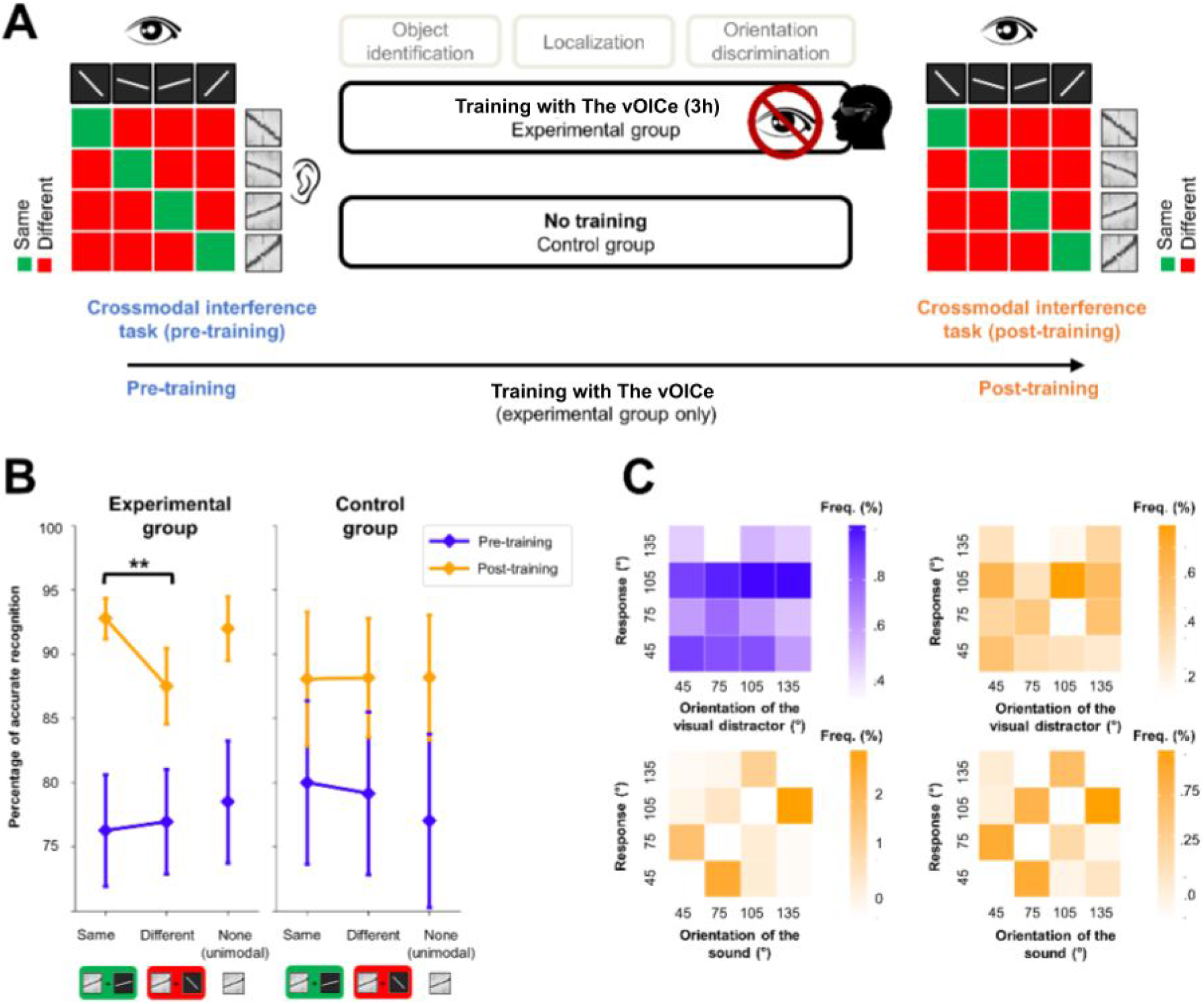
Effect of training with the vOICe on the crossmodal interference task. **A.** Description of the three phases of the experimental protocol. For the experimental group, the crossmodal interference task was performed before (pre-training) and after (post-training) training with the vOICe. For the control group, the crossmodal interference task was performed twice, without training with the vOICe in between. **B.** Mean accuracy (percentage of correct recognition) in the crossmodal interference task across participants of the experimental and control groups, as a function of the Session (pre-training, post-training) and the Type of visual distractor (same, different, none). Error bars represent standard error of the mean. \*\**p* < .01. **C.** Analyses of the distribution of erratic responses made in the crossmodal interference task by the participants involved in the experimental group during the pre-training (in blue) and post-training (in orange) sessions. In the top row, the erratic responses are classified as a function of the orientation of the visual distractor. In the bottom row, the erratic responses are classified as a function of the orientation of the auditory target.

### Crossmodal interference task

In the crossmodal interference task, the participants were asked to recognize a set of 4 soundscapes, which corresponded to the auditory conversion of differently oriented visual lines by The vOICe. Simultaneously, they were presented with visual lines that corresponded to one of the four auditorily converted lines (bimodal block, see Figure 1B). The auditory target and the visual distractor had either the same orientation, or a different one (see Figure 1C). These visual lines are irrelevant to performing the task. There was also a unimodal block in which the auditory targets were presented alone, i.e. without visual distractors.

The auditory targets were generated using the applet The vOICe. They corresponded to differently-oriented white lines on a black background. Four orientations were presented: 45°, 75°, 105°, and 135° relative to the vertical, with a clockwise rotation. The sounds were presented at an intensity of 65 dBA. The visual distractor consisted of oriented white lines on a black background. They were 10° visual angle long, 0.5° thick, and had a contrast of 0.42. The same four orientations as those of the auditory targets were used.

Each trial began with the simultaneous presentation of both a visual and an auditory mask, lasting 1500 ms. Then, a fixation cross was presented at the centre of the screen for 1000 ms, followed by the 1000 ms auditory target and the visual distractor. Trials were separated by a 2000-ms inter-stimulus interval. The participants were asked to press one key, out of four, that corresponded to the sound they heard. The four adjacent keys (F, G, H, and J) on the computer keyboard were used. The participants were instructed to respond with their dominant hand, one finger per key. Four different associations between the keys and the different auditory targets were used and counterbalanced across participants. Emphasis was put on accuracy rather than on speed.

After a short 2-minutes familiarization phase, during which the participants actively learned the mapping between the keys and the auditory targets, they completed two blocks of trials. The unimodal block consisted of 20 training trials with feedback, and 40 test trials without feedback (10 presentations of each auditory target). The bimodal block consisted of 20 training trials with feedback and 80 test trials without feedback (five presentations of each of the 16 possible audio-visual combinations, i.e. 4 auditory targets by 4 visual distractors). The order of the unimodal and bimodal blocks was counterbalanced between participants. Half of participants began with the unimodal block, the other half with the bimodal block.

### Auditory tests

Four tests were used to measure the participants’ auditory abilities, which corresponded to the auditory features relevant for The vOICe, namely: intensity, pitch, duration, and extraction of complex sounds in noisy background. Intensity, pitch, and duration thresholds were measured using a two alternative forced choice. Based on a study by Kidd, Watson & Gygi (2007), two sounds were presented consecutively, a standard one and a deviant one, in a random order. The participants’ task was to indicate which sound corresponded to the standard one. The standard sound was set at 1000 Hz, 65 dBA, and 500 ms. The deviant tone was the same as the standard one, except for the relevant feature (e.g. for the pitch discrimination task, deviant tone is 1000 Hz + ∆ Hz, 65 dBA, 500 ms). The deviation (∆) was adjusted at each trial, based on the participant’s previous answer, using a maximum-likelihood procedure (Green, 1993; Kidd, Watson, Gygi, 2007). This procedure is known to converge rapidly toward the participant’s threshold. Each test was composed of three blocks of 30 trials. As the participants completed these tests twice (once in pre-training and once in post-training session), their final score consisted in the mean threshold of the six blocks.

The capacity to extract complex sounds in noisy background was measured with the HINT (Nilsson, Soli, & Sullivan, 1994, Vaillancourt *et al.*, 2005). This test consists in the presentation of sentences in a speech-corrected white noise background. The participants were instructed to repeat the sentences they heard as accurately as possible. Noise was presented at a fixed level of 65 dBA. The signal-to-noise ratio was adjusted adaptively according to the participants’ previous answer. A first list of 21 sentences was presented as a training list. Then the test included four lists of 21 sentences each. Each list gave a signal-to-noise ratio (SNR), which is the difference between the level of the speech and the level of the noise needed to achieve 50% recognition. The participants’ final SNR was the mean of the SNR calculated for the eight test lists (four in pre-training and four in post-training sessions).

### Training tasks with The vOICe

The training phase with the vOICe was designed as a short version of the one used by Stiles, Zheng, and Shimojo (2015). It was divided into two 1.5-hour sessions, which took place over two different days. Each session included the following tasks, completed in the same order: shape recognition, target localization, and orientation recognition (see Figure 2A).

For the shape recognition task, the participants had to match the soundscapes from The vOICe with the corresponding images. In each trial, the participants heard a soundscape in the headphones and, at the same time, four different images were presented on the screen. The participants had to choose which image corresponded to the soundscape. They could hear the soundscape as many times as they wanted. Fifty-six different sounds were presented four times each, in both sessions, so that each participant had to recognize 448 sounds during the whole training. The complexity of the images and the sounds gradually increased, from simple points to complex geometrical shapes, as reported in several studies using a training phase with a conversion device (e.g., Kim & Zatorre, 2008; Striem-Amit *et al.*, 2012). The participants completed as many trials as they could during each 30-minute session.

For the localization task, the participants were blindfolded and equipped with the device. They wore a pair of 3D-printed glasses with a camera embedded inside them. The visual images recorded by the camera were converted on-line into sounds by The vOICe. The resulting sounds were transmitted to the participants via headphones. The task consisted of pointing toward a 5-cm black circle, placed randomly on a white table by the experimenter. The participants completed as many trials as they could during each 30-minute session.

For the orientation recognition task, the participants were also blindfolded and equipped with the device. Differently-oriented white lines were presented on a black screen. There were six possible orientations that appeared in a random order: 0°, 45°, 67.5°, 90°, 112.5°, and 135°, relative to the vertical, with a clockwise rotation. The participants heard the on-line auditory conversion of these lines through the device the VOICe. Responses were given using six keys of the keyboard (1, 2, 3, 4, 5, and 6). The participants completed as many trials as they could during each 30-minute session.

## Results

### Trained participants resort to visual processes to recognize soundscapes

To analyse the accuracy (0: incorrect, 1: correct) in the sound recognition task, a generalized linear mixed model was used with the method of model comparison (Moscatelli *et al.*, 2012), using Session (pre-training, post-training) and Similarity between the orientations of the auditory and visual stimuli (same, different) as fixed effects, and participants as random effect. The interaction Session × Similarity was significant (p < .05, *β* = −0.67 ± 0.29) (see Figure 2B). Before training, there was no significant difference in accuracy when the sound and the visual distractor had the same orientation (mean 76.3% ± 16.8) or a different one (76.9% ± 15.8) (p = .80, *β* = 0.04 ± 0.16). However, after training with the device, accuracy in sound recognition was significantly higher when the visual distractor was similar to the sound (92.7% ± 6.2) than when it was different (87.5% ± 11.4) (p < .01, *β* = −0.65 ± 0.24). Thus, there was a visual interference effect on the auditory task only after training with the visual-to-auditory conversion device. The same analysis was conducted on response time (RT) data, and showed a significant effect of Session (p < .001, *β* = −138.98 ± 24.99), but no effect of Similarity (p = .18, *β* = −105.86 ± 79.10), and no interaction between these two factors (p = .68, *β* = 20.81 ± 50.00). Thus, the visual interference effect appeared for accuracy only. It is to be noted that, in the instructions given to the participants, the stress was laid on accuracy, not on response latencies, which can explain that the effect was obtained only in the former case.

In order to assess whether the interference effect consists in facilitating the same trials or disturbing the different ones, a unimodal block was run, either just before or just after the bimodal block. In this unimodal block, the sounds were presented without any visual distractor. In the pre-training session, there was no significant differences between the unimodal (78.5% ± 18.4) and the bimodal same (p = .40, *β* = 0.15 ± 0.18) conditions, nor between the unimodal and the bimodal different conditions (p = .44, *β* = 0.10 ± 0.13). In the post-training session, there was no significant difference between the unimodal (92.0% ± 9.6) and the bimodal same conditions (p = .68, *β* = −0.11 ± 0.26). However, the accuracy was significantly higher in the unimodal than in the bimodal different condition (p < .005, *β* = 0.55 ± 0.18). Thus, the interference effect is characterized by a disturbance effect rather than by a facilitation one.

In order to verify whether the visual interference effect might be due to the mere repetition of the sound recognition task, participants of a control group completed the crossmodal interference task twice, without being trained with the device between the first and the second sessions (see Figure 2B). Two results were expected: an effect of task repetition and a lack of effect of the crossmodal interference. This corresponds to what was observed. The obtained results showed in the control group (as in the experimental group) an effect of task repetition, with overall higher performance in the second session (78.9% ± 25.4) than in the first one (88.1% ± 19.2, p < 10^−9^, β = 0.96 ± 0.15). On the other hand, no difference was found between the conditions same and different (first session, 80.1% ± 25.5 versus 79.2% ± 25.3, p = .68, β = 0.08 ± 0.20; second session 88.1% ± 21.1 versus 88.1% ± 18.5, p = .99, β = 0.00 ± 0.24) as well as no Session × Similarity interaction (p = .80, β = −0.07 ± 0.30). This confirms that a mere repetition of the task cannot induce an interference effect. However, the three way interaction did not reach significance (p = .12, *β* = −0.66 ± 0.42). This can be accounted for first by the fact that a visual interference effect was expected in one condition (i.e. the second session of the experimental group) compared to a lack of effect in the remaining three conditions (i.e., in the first session of both groups and in the second session of the control group). Second, a strong repetition effect in both groups (experimental group, p < 10^−9^, β = 0.79 ± 0.13, control group, p < 10^−9^, β = 0.96 ± 0.15) captures most part of the variance thereby hiding the smaller visual interference effect. Consequently, obtaining a three way interaction would have required a very large sample size (simulation indicates an N > 60 per group for a power of 80%). Thus, the visual interference effect was obtained only in the expected condition (experimental post-training session 92.7% ± 6.2 versus 87.5% ± 11.4, p < .01, β = −0.65 ± 0.24; experimental group first session 76.3% ± 16.8 versus 76.9% ± 15.8, p = .80, β = 0.04 ± 0.16; control group first session, 74.1% ± 31.3 versus 74.6% ± 28.7, p = .82, β = 0.04 ± 0.20; control group second session 82.0% ± 27.1 versus 82.7% ± 24.5, p = .91, β = 0.03 ± 0.24, as illustrated in Figure 2B). In summary, the results can be taken as illustrating that processes shared with vision were involved only in case of proper learning with a sensory substitution device.

In order to ensure that the interference effect does not arise from a misunderstanding of the instructions, with the participants responding to the visual distractor instead of the auditory target, error distribution was analysed. The contingency tables were computed for the pre-training and post-training sessions, across the participants involved in the experimental group only (see Figure 2C). Classifying erratic responses according to the visual distractor shows a contingency table that is not biased either in the pre-training (χ^2^ = 4.07, p = 0.907) or in the post-training session (χ^2^ = 13.7, p = 0.134). Conversely, classifying erratic responses according to the auditory target clearly shows a biased table, both in the pre-training (χ^2^ = 426.0, p < 0.001) and in the post-training (χ^2^ = 150.0, p < 0.001) sessions. When a visual distractor with a different orientation than that of the auditory target is presented, the participants do not confuse the auditory and visual orientations, nor do they mistake visual orientations for auditory ones. These results are consistent with the idea that, even when the participants make mistakes, they still follow the instructions and try to recognize the auditory target.

### Controlling for crossmodal correspondence effects

Given the features of the auditory and visual stimuli that were used, and the nature of the interference task, one could argue that the obtained interference effect merely reflects the existence of crossmodal correspondences. Indeed, it is a well-known fact that natural associations occur between different sensory modalities, *e.g.*, between auditory pitch and visual or tactile elevation, or between auditory loudness and visual brightness (Deroy *et al.*, 2016; Spence, 2011, for a review). These crossmodal correspondences have been shown to influence participants’ responses in tasks that are similar to the crossmodal interference task used in the present study (Evans & Treisman, 2009). Given that the pitch of ascending and descending sounds can be naturally associated, respectively, with ascending and descending visual lines, crossmodal correspondences might have played a role in the crossmodal interference task.

In order to evaluate this possible role of crossmodal correspondences, the results of the participants involved in the experimental group were analysed as a function of the congruency between auditory and visual stimuli. Trials in which both the auditory target and visual distractor were ascending or descending were labelled as “congruent”. Trials in which the auditory target and visual distractor differed were labelled as “incongruent”. A generalized linear mixed model was used to analyse the accuracy (0: incorrect, 1: correct) in the crossmodal interference task, as a function of Session (pre-training, post-training) and Congruency (congruent, incongruent) as within-participant factors. There is no effect of Congruency in the pre-training (*p* = 0.43, *β* = 0.11 ± 0.14), nor in the post-training (*p* = 0.13, *β* = −0.28 ± 0.19) sessions (see Figure 3).

**Figure 3.**
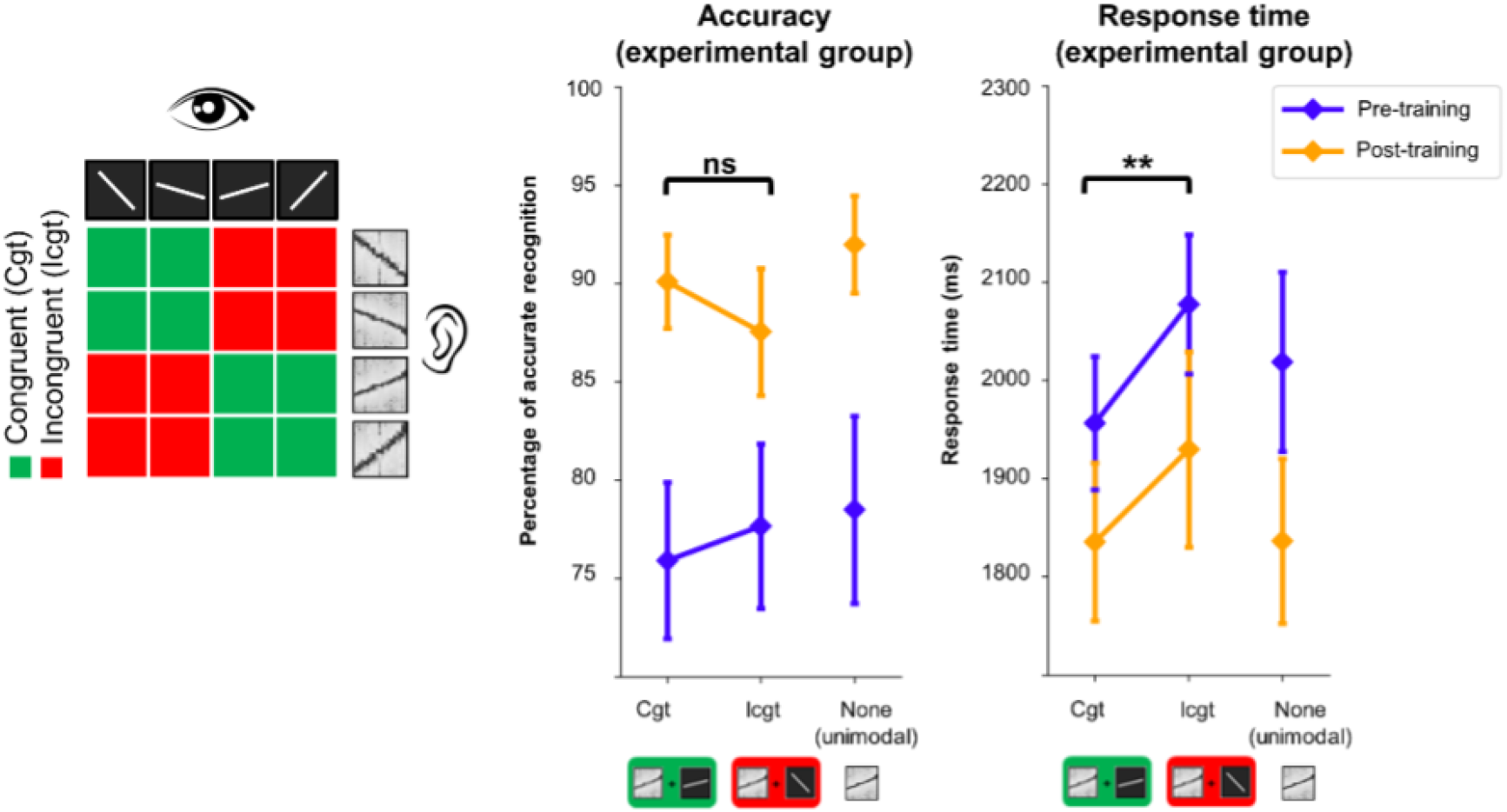
Control analyses of the participants’ responses for congruency effects. On the left: Description of congruent and incongruent trials. In congruent trials, the auditory target and visual distractor were both ascending or descending, independently of the slope. In incongruent trials, one was ascending and the other one descending. On the right: Accuracy (percentage of correct recognition) and response times obtained by the participants involved in the experimental group in the crossmodal interference task as a function of Session (pre-training, post-training) and Congruency (congruent, incongruent, none) between the orientation of the auditory target and that of the visual distractor. None refers to the unimodal condition.

The same analysis on RT data showed a main effect of Congruency (p < 0.05, *β* = 149 ± 68.3). However, the Session × Congruency interaction was not significant (p = 0.53, *β* = −26.8 ± 43.1). This suggests that Congruency has the same effect in the pre-training (p < 0.001, *β* = 122 ± 31.0) and in the post-training (p < 0.001, *β* = 95.1 ± 28.4) sessions (see Figure 3). Overall, the analyses conducted on congruency revealed the effect of crossmodal correspondences on response times, but not on accuracy, with faster responses when the auditory target and the visual distractor were both ascending or descending than when they were headed in opposite directions. However, the effect of congruency was not influenced by training with the device. Thus, the visual interference effect that appeared after training with the device results from a change in the involved processes, and cannot be explained by the mere existence of natural associations between the characteristics of the auditory targets and visual distractors.

### The processes involved are derived from the auditory modality

During the use of the visual-to-auditory conversion device, even though visual processes are involved, audition may still play a role in the processing of the input signal. This was investigated in our study by taking into account the participants’ auditory abilities. To do so, these abilities were measured by means of discrimination tasks of pitch, loudness, and duration of sounds, which correspond to the three auditory features relevant to the device’s use, and with an additional task evaluating the participants’ abilities to extract complex sounds in a noisy background. For each participant, a global auditory score was computed and subsequently correlated with the individual performance they obtained during both the sound recognition task and the training phase with the device (see the Material and Methods section for the description of the device training score). The auditory score was found to correlate with the global accuracy in the auditory task (*r* = .60; *p* < .05), but not with the visual interference effect (*r* = .22; *p* = .42). These results show that the extent to which participants spontaneously resort to visual images when hearing the device’s sounds does not depend on their individual auditory abilities. However, individual auditory abilities were found to influence the use of the device during the training phase. The training scores correlated significantly with the auditory scores (*r* = .70; *p* < .01): the higher the auditory abilities, the higher the performance with the device. Therefore, to summarize, learning to use a conversion device relies not only on the ability to visualize sounds, but also on the ability to process the auditory input signal.

### Sensory phenomenology varies as a function of the task and individual auditory abilities

In order to further characterize the nature of the involved processes, the associated phenomenology was evaluated (see Table 1). The participants completed questionnaires about their subjective experience with the device during the training phase. They were asked to rate on 5-point Lickert scales (1-low to 5-high) the extent to which their experience with the device resembled audition, vision, and a sonar-like experience. The phenomenology was unsurprisingly mainly auditory (4.8/5, SD = 0.4). However, visual (3.5/5 ± 0.6) and sonar-like (3.6/5 ± 0.9) phenomenologies were also reported. The reported phenomenologies varied as a function of the task, as was previously observed (Auvray *et al.*, 2007), and as a function of participants’ auditory abilities. For those participants with high auditory abilities (median split), the phenomenology was more visual in the object recognition task (4.8/5 ± 0.5) than in the localization task (2.9/5 ± 0.5) (*t*(7) = 4.09, *p* < .01), whereas it was more sonar-like in the localization task (4.0/5 ± 1.0) than in the object recognition task (2.7/5 ± 1.3) (*t*(7) = 2.97, *p* < .05). For those participants with low auditory abilities, the reported phenomenology did not differ as a function of the task for the visual phenomenology (3.2/5 ± 0.4, 2.4/5 ± 0.8; *t*(7) = 1.54, *p* = .17) or for the sonar-like one either (3.9/5 ± 0.9, 3.4/5 ± 0.9; *t*(7) = 1.32, *p* = .23). Altogether, the phenomenological reports indicate that high auditory abilities allow people to extract the most relevant sensory information for a given task, resulting in different sensory experiences across tasks. In particular, in object recognition, shape is the most relevant information, whereas in localization, it is distance. When focusing on distance, participants may be biased toward elaborating a sensory experience similar to using sonar or echolocation devices, which consists of computing distance from time information, a very frequent process in animals (Halfwerk *et al.*, 2014). By contrast, low auditory abilities may prevent people from reaching a differentiated use of the device, associated with a differentiated phenomenology.

**Table 1.** Phenomenologies associated with The vOICe’s use as a function of the task and session, averaged across participants in the experimental group. Pre-training and post-training refers to questionnaires administered each time just after the interference task. Note that, as the sonar-like experience often reappeared in participants’ verbal reports (*e.g.*, see Auvray *et al.* 2007) we added this scale in the questionnaire.

|  | Auditory | Visual | Tactile | Gustatory | Olfactory | Sonar-like |
| --- | --- | --- | --- | --- | --- | --- |
| **Pre-training** | 4.9 | 3.4 | 1.3 | 1.1 | 1.1 | 3.3 |
| **Training 1: recognition** | 4.8 | 4.1 | 1.2 | 1.0 | 1.0 | 3.4 |
| **Training 1: localisation** | 4.8 | 2.6 | 1.6 | 1.1 | 1.0 | 3.9 |
| **Training 1: orientation** | 4.8 | 2.6 | 1.3 | 1.1 | 1.1 | 3.4 |
| **Training 1: mean** | 4.8 | 3.1 | 1.4 | 1.0 | 1.0 | 3.6 |
| **Training 2: recognition** | 4.9 | 4.3 | 1.1 | 1.1 | 1.1 | 3.1 |
| **Training 2: localisation** | 4.7 | 2.8 | 2.0 | 1.1 | 1.0 | 4.1 |
| **Training 2: orientation** | 4.9 | 2.9 | 1.5 | 1.0 | 1.0 | 2.9 |
| **Training 2: mean** | 4.8 | 3.3 | 1.5 | 1.0 | 1.0 | 3.4 |
| **Post-training** | 4.8 | 3.9 | 1.4 | 1.0 | 1.0 | 3.3 |

## Discussion

Our study highlights that using a device that changes the way we receive the inputs from our environment modifies how the brain processes these stimuli. First, the results from the crossmodal interference task showed that, before training, the visual images did not influence the participants’ responses. After training, however, they interfered with the auditory recognition task. In particular, they disturbed the participants’ responses when the auditory soundscape did not correspond to the conversion of the visual image. This visual interference effect reveals a rapid functional plasticity, as users, once trained, cannot prevent themselves from processing the auditory soundscapes with processes that are shared with vision. Second, the participants’ individual auditory abilities also played a role. The correlations between the participants’ auditory scores and their performance with the device suggest that those participants with higher auditory abilities performed better during the training tasks with the device than those with lower abilities. Third, the participants did not report a unisensory phenomenological experience, but considered it to involve visual, auditory, and sonar-like aspects. In addition, participants with higher auditory abilities made a richer use of the device, as their associated phenomenology changed with the type of task, which was not the case for those with lower auditory abilities. Overall, these results underline that a multisensory processing in sensory substitution occurs at the behavioural and phenomenological levels. These two levels are discussed in link with the vertical integration hypothesis (Arnold *et al.*, 2017) and closely related metamodal/ supramodal view of sensory substitution processing (Amedi *et al.*, 2017; Proulx *et al.*, 2014, 2016).

These two hypotheses predict that, using a sensory substitution device involves processes that are not bound uniquely to the visual nor the auditory modality. Our study shows the involvement of these two modalities. At the behavioural level, the results from the visual interference task suggests that, after training, the auditory stimuli coming from the device are no longer processed exclusively through the auditory pathways. Indeed, if the involved processes were only auditory, the simultaneous presentation of visual images should not have interfered with the auditory task. On the contrary, the visual interference effect obtained here suggests that, when hearing sounds, trained participants spontaneously resort to visual images. As a consequence, they are disturbed when the auditory stimulus and the simultaneous visual distractor have a different orientation. It should be noted that the question of whether the interference appears at the visual or at a supramodal level remains open. Nonetheless, our results clearly demonstrate that training with a visual-to-auditory conversion device involves additional processes, different from auditory processes. Second, auditory processes are involved as well. Indeed, participants’ performance with the device depends on their individual auditory abilities, as shown by the correlations between the participants’ individual auditory scores and their performance with the device. Thus, the visual-to-auditory stimuli appear to be processed in an integrated way, which takes into account both the initial features of the stimuli (here auditory), and those novel ones that arise when using the device (here visual). This line of reasoning extends beyond the field of sensory substitution, as an increasing number of scientists defend the view of a brain organized in a task-specific and modality-independent architecture (Pascual-Leone & Hamilton, 2001, Heimler, Striem-Amit, & Amedi, 2015). More generally, our results reveal that sensory plasticity in humans is a complex phenomenon which depends both on the kind of processes that are involved and on individual specificities.

The vertical integration hypothesis also posits that using a conversion device involves flexible processes and phenomenologies, with different weights attributed to the sensory modalities as a function of individual differences and as a function of the task demands. Our results show that different phenomenologies are indeed involved as a function of the task performed with the device – i.e., visual for an identification task, sonar-like for a localization task –, but only for participants with high auditory abilities. This suggests that a differentiated use of the device requires high individual capacities in processing the input signal. Thus, learning to use a conversion device could involve different perceptual strategies, as a function of these pre-existing capacities.

The last point has implications in the field of the rehabilitation of visual impairments through the use of conversion devices. Given that these devices allow their users to compensate for the loss of one sensory modality, the most optimal way to learn how to use these devices is crucial. Our results suggest that an optimal learning procedure will consist in training participants both to accurately analyse the features of the stimuli provided to the initial sensory modality, and to correctly interpret its translation into the other sensory modality. In addition, the balance between these two learning components should be adjusted as a function of users’ individual differences concerning low-level and high-level perceptual abilities. More broadly, our study has an impact on the understanding of how people adapt to a new environment or to a new tool, namely by relying on individual low level and high level abilities and strategies.

## Conclusion

To conclude, William James made the hypothesis that, if our eyes were connected to the auditory brain areas, and our ears to the visual brain areas, we would “hear the lightning and see the thunder” (James, 1890). The results of our study reveal that using a device that changes the way we receive inputs from the environment simultaneously changes the way these stimuli are processed. Thus, the stimuli are not only processed by different brain networks, they are also processed differently at the functional and phenomenological levels. However, the association of the visual interference effect with the role of individual auditory abilities and the associated phenomenology underlines the fact that functional plasticity is complex, and based on a multisensory architecture involving both visual and auditory processes. Our results show that people can become able to visualize auditory stimuli while keeping on processing it auditorily. In William James’ words, they become able to see the thunder while keeping on hearing it.

## Author contributions

J.P.-L., G.A., and M.A. designed the experiments and wrote the paper. J.P.-L. conducted the experiments. J.P.-L. and G.A. performed the statistical analyses.

## Acknowledgements

This work was supported by the Labex SMART (ANR-11-LABX-65) and the Mission pour l’Interdisciplinarité (CNRS, Auton, Sublima Grant). We thank Maxime Ambard for the glasses used in this experiment, Sylvain Hanneton for the useful methodological and statistical discussions, and Xavier Job for his comments on the manuscript. GA was funded by a grant from the European Research Council (Wearhap, FP7, N°601165).

